# Object motion influences feedforward motor responses during mechanical stopping of virtual projectiles: A preliminary study

**DOI:** 10.1101/2021.04.20.440704

**Authors:** Ana Gómez-Granados, Isaac Kurtzer, Sean Gordon, Deborah A. Barany, Tarkeshwar Singh

**Author notes:** share senior authorship (Deborah A. Barany, Tarkeshwar Singh). Corresponding author: Tarkeshwar Singh, 32 Recreation Building, The Pennsylvania State University, University Park, PA-16802.

## Abstract

An important window into sensorimotor function is how humans interact and stop moving projectiles, such as stopping a door from closing shut or catching a ball. Previous studies have suggested that humans time the initiation and modulate the amplitude of their muscle activity based on the momentum of the approaching object. However, real-world experiments are constrained by laws of mechanics, which cannot be manipulated experimentally to probe the mechanisms of sensorimotor control and learning. An augmented-reality variant of such tasks allows for experimental manipulation of the relationship between motion and force to obtain novel insights into how the nervous system prepares motor responses to interact with moving stimuli. Existing paradigms for studying interactions with moving projectiles use massless object and are primarily focused on quantifying gaze and hand kinematics. Here, we developed a novel collision paradigm using a robotic manipulandum where participants mechanically stopped a virtual object moving in the horizontal plane. On each block of trials, we varied the virtual object’s momentum by increasing either its speed or mass. Participants stopped the object by applying a force impulse that matched the object momentum. We observed that arm force increased as a function of object momentum linked to changes in virtual mass or speed, mirroring results from studies involving catching free-falling objects. In addition, increasing object speed resulted in later onset of hand force relative to the impending time-to-contact. These findings show that the present paradigm can be used to determine how humans process projectile motion for hand motor control.

## Introduction

An important visuomotor task that humans perform with relative ease is stopping projectiles. For example, in everyday contexts humans may be tasked with stopping a door with a spring from closing shut or catching a saltshaker slid across the table. Similarly, many sports rely on stopping moving objects, such as a hockey goalie stopping a puck with their stick or a soccer player stopping the ball with their foot. These actions require that the activity of antagonist limb muscles be appropriately scaled and precisely timed to absorb the projectile momentum during contact.

In a series of seminal studies, Lacquaniti and colleagues showed that projectile kinematics directly influenced activation of muscles both before and during impact (Lacquaniti and Maioli 1989b; Lacquaniti and Maioli 1989a; Lacquaniti et al. 1991; Lacquaniti et al. 1992; Lacquaniti et al. 1993). They showed two interesting results. First, the anticipatory muscle responses were tuned parametrically to the estimated momentum (mass x speed) of the projectile (reviewed in Fig. 3, Zago and Lacquaniti 2005) such that the consequent increase in limb impedance stabilized the arm during the large and transient transfer of momentum from the projectile to the arm. Second, projectile speed appeared to have little to no effect on the timing of anticipatory muscle activation prior to contact (reviewed in Zago and Lacquaniti 2005). However, since these studies primarily used balls falling freely under gravity, they could not manipulate the motion of the projectiles to test how motion-processing of moving projectiles contributes to timing and scaling of limb motor responses.

The goal of the present study is to understand how the visuomotor system transforms motion signals to control interactions between the body and moving objects. To that end, we developed an augmented-reality based paradigm. We simulated the physics of mechanical interactions between the hand and a projectile moving in the horizontal plane (no acceleration). We call this the *Mechanical STopping Of virtual Projectiles (MSTOP)* paradigm. This paradigm allows us to replicate the mechanical interaction between a projectile and the hand when humans apply limb force to stop moving projectiles. In our paradigm, we programmed a robotic manipulandum to replicate the mechanical interaction between a projectile and the hand during collision. We did that by simulating the physical equivalence between momentum and applied impulse (area under force-time curve) defined by Newton’s Second Law. We also adopted an explicit and strict task criterion - to successfully stop the projectile, participants must apply a force impulse within a 5% margin of error to the momentum of the projectile. Our paradigm is different from existing paradigms that use massless virtual objects to quantify how humans prepare and execute interception movements to moving objects (Tresilian 2005; Mrotek and Soechting 2007; Brenner and Smeets 2011). Specifically, our paradigm also allows us to measure how humans control limb force and stabilize posture during interactions with moving projectiles.

In this paper, we introduce this new motor control paradigm with a simple experiment. We tested how systematically varying the momentum of a projectile by changing either its speed or its mass affected limb force. We varied the mass and speed of virtual objects moving towards the participant in different blocks and measured limb forces before and during contact between the hand and the virtual object. Based on previous reports (reviewed in Zago and Lacquaniti 2005), we predicted that the amplitude of the feedforward limb forces prior to the contact would scale with object momentum whether the momentum increased due to speed or mass. Our results confirmed this expectation; participants increased limb force prior to impact based on the momentum of the object. In addition, our virtual paradigm allowed us to decouple the mass and speed of the object to test the prediction that motor response timing is based on a continuous estimate of time-to-collision (Lee 1976; Tresilian 1991). In contrast to reports on catching free-falling objects that found timing is invariant to object motion characteristics (Lacquaniti and Maioli 1989b), we observed that participants increased limb force closer to the time of impact at higher momentums, especially when the increase in object momentum was due to an increase in its speed. These proof-of-concept results will allow us to test specific hypotheses in the future to probe how humans control limb forces in anticipation of a mechanical interaction with moving projectiles.

## Methods

### Participants

Twenty participants (20.6 ± 2.04 years; 10 ♀) completed the study. All participants were right-handed, had no history of neurological disorders, and had no current injuries or pain of the upper limbs. Each participant provided written informed consent prior to participating and were compensated for their time. All procedures were approved by the local Institutional Review Board of the University of Georgia, Athens, GA.

### Apparatus

The task was performed on a robotic manipulandum (KINARM End-Point Lab, KINARM, Kingston, ON, Canada) that participants grasped with their right hand. The robotic arm could be moved in a horizontal plane and a mobile arm support (SaeboMAS, Saebo Inc., Charlotte, NC, USA) was used to support the weight of the participant’s arm to avoid fatigue. Visual objects were generated using an online Gabor-patch generator (https://www.cogsci.nl/gabor-generator), with fixed parameters (orientation: 90°, size: 96 pixels, Gaussian envelope, standard deviation: 24 pixels, frequency: 0.1 cycles/pixel, and phase: 0 cycles). The color scheme of the object was used to differentiate the object’s assigned virtual mass (see below). Note that unlike true Gabor patches, the object had a clear boundary and was easily detectable in the workspace (see Fig. 1).

**Figure 1.**
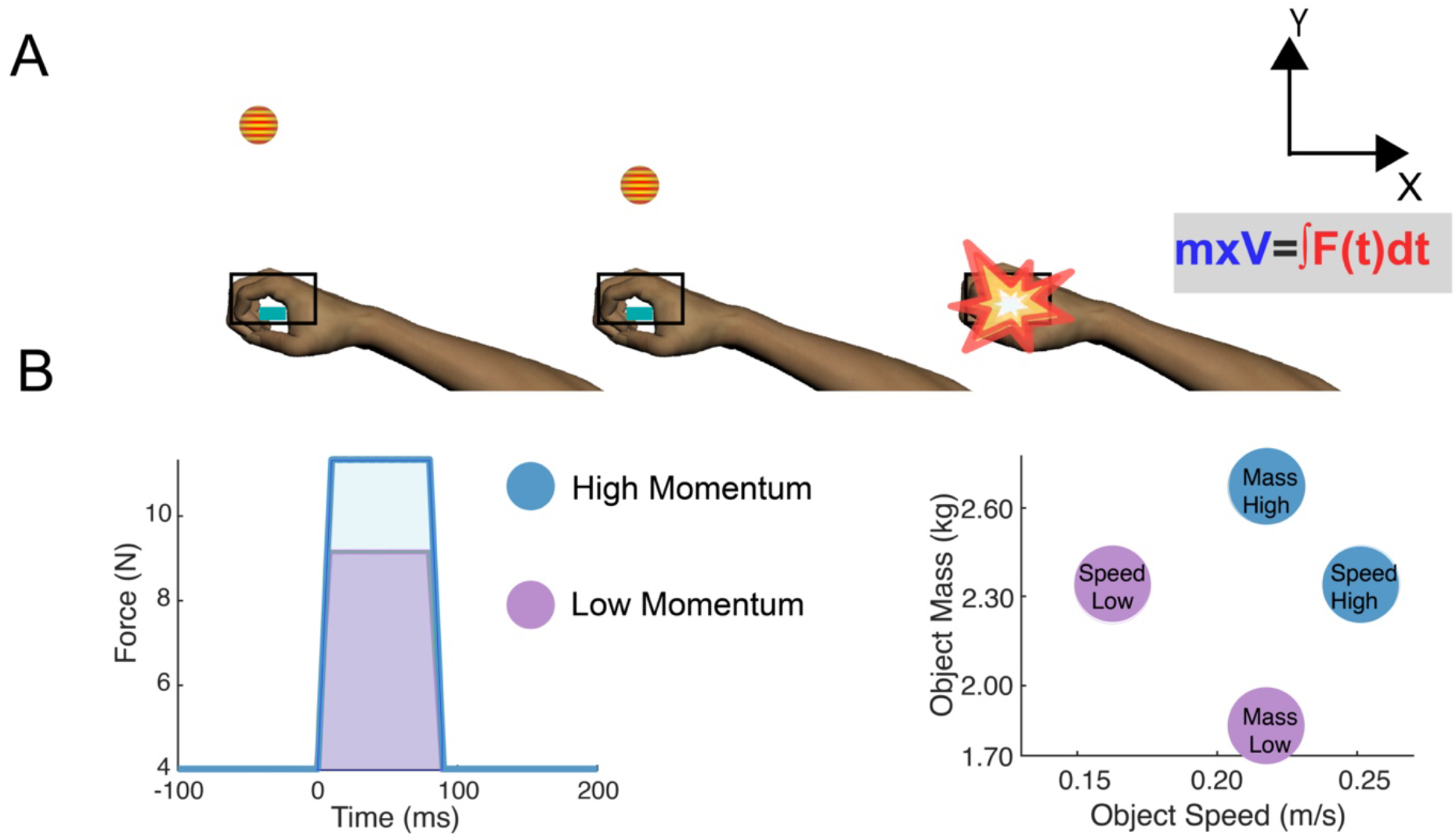
Experimental setup. A) Once participants moved their hand into the rectangular box and stabilized their arm against the background load, a fixation cross appeared (not shown). Participants had to fixate on the cross until it disappeared (600 ms). 200 ms after a circular object appeared at the same position and moved towards the participant. Participants were instructed to match the force applied by the robot during the contact. B) Force-time curve for the perturbation applied by the robot. The area under the curve, the impulse, is exactly equal to the momentum carried by the virtual object. The right panel shows the mass and speed values assigned to each sub-condition.

### Task Design

Participants performed a *Mechanical STopping Of Projectiles (MSTOP*) task where the objective was to “stop” a virtual circular object moving in the negative Y direction (towards the body, Fig. 1A) by applying a force pulse to bring object momentum to 0. We assigned a virtual mass to an object and then multiplied it with its speed (in the KINARM frame of reference) to obtain a momentum for the virtual object. Since the change in momentum and applied impulse (area under force-time curve) are equivalent (Eq. 1, Newton’s Second Law), we converted the momentum of the object into an impulse template (see Fig. 1B) that the KINARM robot applied to the participant’s hand when the object contacted the hand (right panel, Fig. 1A). This allowed us to replicate critical aspects of mechanics of catching.

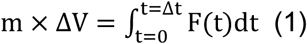

Here, the term on the left-hand side is the change in momentum; the term on the right-hand side is the impulse, the integral of force and time. To bring an object approaching the body with a speed V and an assigned mass ‘m’ to a complete stop, the hand will have to apply a time-varying force F(t) over Δt s. We fixed Δt to 90 ms. When the object contacted the hand, the robot applied the impulse on the lefthand side of Eq. 1 over 90 ms with a 10 ms long rise time and fall time and a 70 ms steady state force. Participants experienced that impulse as simulating a contact between the object and the hand. During the contact between the hand and the object, participants were instructed to try to match that impulse applied by the robot as closely as possible (±5% of the impulse applied by the robot) to bring the object to a stop. Exemplar force profiles applied by the robot are shown in Figure 1B (left panel) for objects with different speeds and masses.

Object momentum (*Low* or *High*) was varied across experimental blocks via changes to either the object’s speed or mass (right panel of Fig. 1B and Table 1). The momentum difference between the *Low* (0.41 kg.m/s) and *High* (0.58 kg.m/s) momentum conditions was ~41%. For each condition, there were two sub-conditions: one in which the difference between *Low* and *High* momentum was due to object speed (*Speed Low* vs. *Speed High*) and one in which the momentum difference was due to object ‘mass’ *(Mass Low* vs. *Mass High*). Object mass was visually represented by using different colors for *Mass High* (dark blue with gray lines on the Gabor patch) and *Mass Low* (light blue with gray lines). In the Speed sub-conditions, the object was always red with yellow lines to denote no changes in mass. The purpose of the sub-conditions was to explore whether any differences in limb motor control observed with increases in momentum depended on specific features of the object dynamics. Importantly, this decoupling of speed and mass can only be achieved in augmented-reality environments.

**Table 1.** Object mass, speed, and momentum/robot impulse values for the experimental sub-conditions.

| Sub-condition | Mass (kg) | Speed (m/s) | Momentum (kg.m/s)<br>or Impulse (N.s) |
| --- | --- | --- | --- |
| **Speed Low** | 2.32 | 0.18 | 0.41 |
| **Speed High** | 2.32 | 0.25 | 0.58 |
| **Mass Low** | 1.87 | 0.22 | 0.41 |
| **Mass High** | 2.65 | 0.22 | 0.58 |

### Procedure

At the beginning of each trial, participants moved the cursor representing their hand location into a designated collision area (5 x 10 cm, rectangle) and maintained a static hold until the end of the trial. If their hand escaped the rectangle, the trial was aborted and repeated. When participants entered their hand in the rectangle, a background load (4 N in the -Y direction, see Fig. 1A) was applied and stayed on for the remainder of the trial. Background loads were applied to stabilize the hand prior to the presentation of the moving object (Singh et al. 2017; Barany et al. 2020) and to minimize unnecessary anticipatory hand movements that would affect the measurement of force onset. After a fixed delay of 1,700 ms, a fixation cross appeared in the middle of the screen (20 cm away from the center of the rectangle). Participants were instructed to fixate on the cross until it disappeared (600 ms). Then the circular object (1 cm radius) appeared on the same position 200 ms after the cross disappeared and immediately started moving in the -Y direction towards the middle of the collision area (Fig. 1A).

Participants were asked to stop the object by matching the force impulse applied by the robot. The precise instruction given to the participant was “match the force applied to the hand so that the object stops.” Participants were allowed to move their hand as long as the hand stayed within the rectangle. It is important to note that the mechanical interaction during the contact is primarily determined by feedforward commands to muscles and muscle’s intrinsic viscoelastic properties (Burdet et al. 2013) because the 90 ms contact duration is too short for voluntary feedback correction. Though fast feedback corrections are observed in muscle activity within the first 50-100 ms after mechanical perturbations (reviewed in Scott 2012; Kurtzer 2015), hand force is likely low-pass filtered by the musculoskeletal system (Burkholder 2016) and consequently may not include effects of feedback responses.

The performance of a participant in three different trials of the Speed Low sub-condition along with the impulse applied by the robot (Robot Force as a function of time) is shown in Figure 2A. For ideal task performance, the limb force should mirror the robot force. However, ideal performance is almost never achieved because Newton’s Third Law does not apply to a free limb, i.e., a limb is not a rigid body. Other groups have also modeled non-rigid mechanical interactions during collision between the hand and a moving projectile using Hooke’s Law (Kuling et al. 2019). Thus, because the two reactive forces (object on hand and hand on object) are not equal, we provided a ±5% force error margin for participants to be successful on a trial. Note that the 5% error margin was a fairly strict criteria as the Weber fraction for force production is between 10-20% (Jones 1986; Debats et al. 2012).

**Figure 2.**
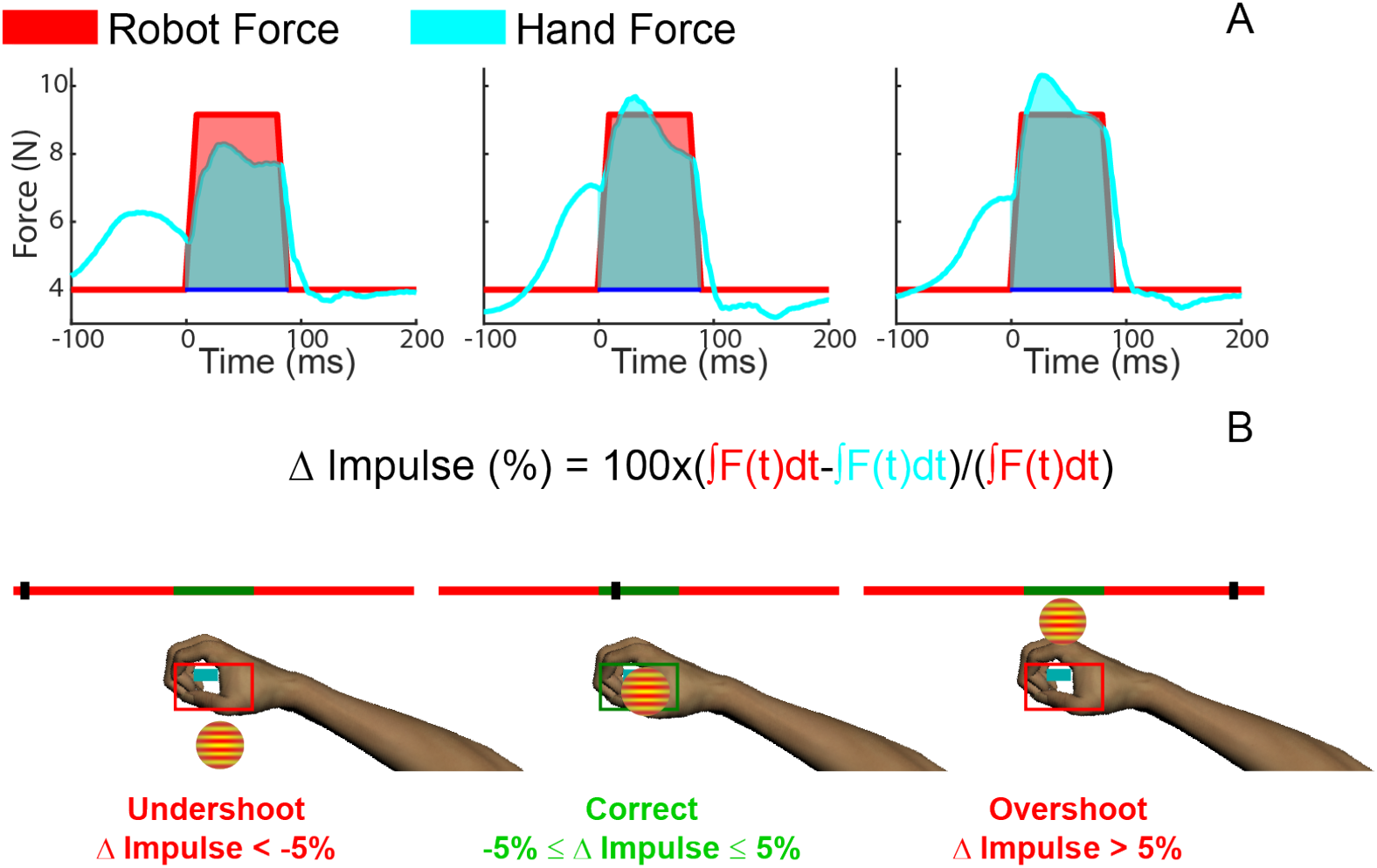
Performance feedback. A) Exemplar trials from a participant in three different situations when the participant applied less than 95% of the required impulse (left, undershoot), an impulse within 95-105% of the desired impulse (middle, correct), and more than 105% of the impulse (right, overshoot). The red shaded area is the impulse applied by the robot and the cyan area is the impulse applied by the participant. The red and cyan curves are the force applied by the robot and the hand, respectively. B) The corresponding feedback provided to participants for the three trials in A. The black bar on the horizontal line indicated how far performance for that trial deviated from the ideal impulse (middle of green section of the horizontal line).

Participants were given two forms of feedback – one with the object and another one with a visual scale (see Fig. 2B). Object feedback had three different levels: if the impulse was matched within a ±5% margin of error (see Eq. 2), that would imply that an appropriate impulse was applied to match the momentum of the object and the object would stop after the contact (reflecting a successful stop). If the impulse applied by the participant was higher than the 5% margin of error, that would imply that more than the necessary impulse was applied, and the object would “fly back” and move in the +Y direction (overshoot). Lastly, if the impulse applied by the participant was lower than the margin of error, the object would continue unabated on its original trajectory in the -Y direction towards the participant (undershoot).

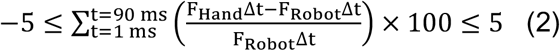

In addition to discrete object feedback, participants were provided a more continuous measure of performance by adding a visual analog scale that had a green center region surrounded by red region. The scale was set up based on Eq. 2, with the green region indicating the ±5% margin of error and the red regions indicating how low or high the applied impulse was from the impulse required to bring the object to rest. Feedback was displayed for 1,700 ms, followed by an inter-trial delay of 2,500 ms.

Participants performed two blocks of 35 trials for each of the four sub-conditions (8 blocks, 280 trials total). The block order was counterbalanced across participants such that half of the participants performed the four Mass sub-condition blocks first and the other half performed the four Speed subcondition blocks first. Within each sub-condition, the block order (e.g., Momentum *High* or *Low*) was randomized for each participant and across participants.

### Data recording and analysis

All analyses were performed in MATLAB (Mathworks R2020b, Natick, MA). We treated the first three trials in each block as practice and only performed analyses on the remaining 32 trials. Hand kinetics and kinematics were sampled at 1 kHz and digitally low-pass filtered (second-order, dual-pass Butterworth, 50 Hz effective cutoff). For each sub-condition, we calculated the number of trials in which the hand impulse (1) successfully matched the robot impulse within the 5% margin of error (correct), (2) was above the margin of error of the robot impulse (overshoot), and (3) was below the margin of error of the robot impulse (undershoot). For each trial within a block, we calculated the overall hand impulse applied as well as the percentage difference between the impulse applied by the robot and the participant (see Equation 2), ΔImpulse (%).

Participants typically increased their hand force and moved their hand slightly toward the object (1-3 cm) right before contact. Thus, we quantified when the hand force and hand speed increased above baseline values in anticipation of the contact with the object during the static hold. Since analyses based on hand speed produced similar results to analyses of hand force, we used the force data for all subsequent analyses.

We first calculated the mean force applied against the background load in an interval (t = [−500, −400 ms]) prior to the contact and then subtracted it from the peak force recorded from the force sensor during contact. Then starting from the point of contact (t = 0), we went backward in time and defined hand force onset as the first time point when the hand force dropped below 5% of the peak force value (Fig. 3).

**Figure 3.**
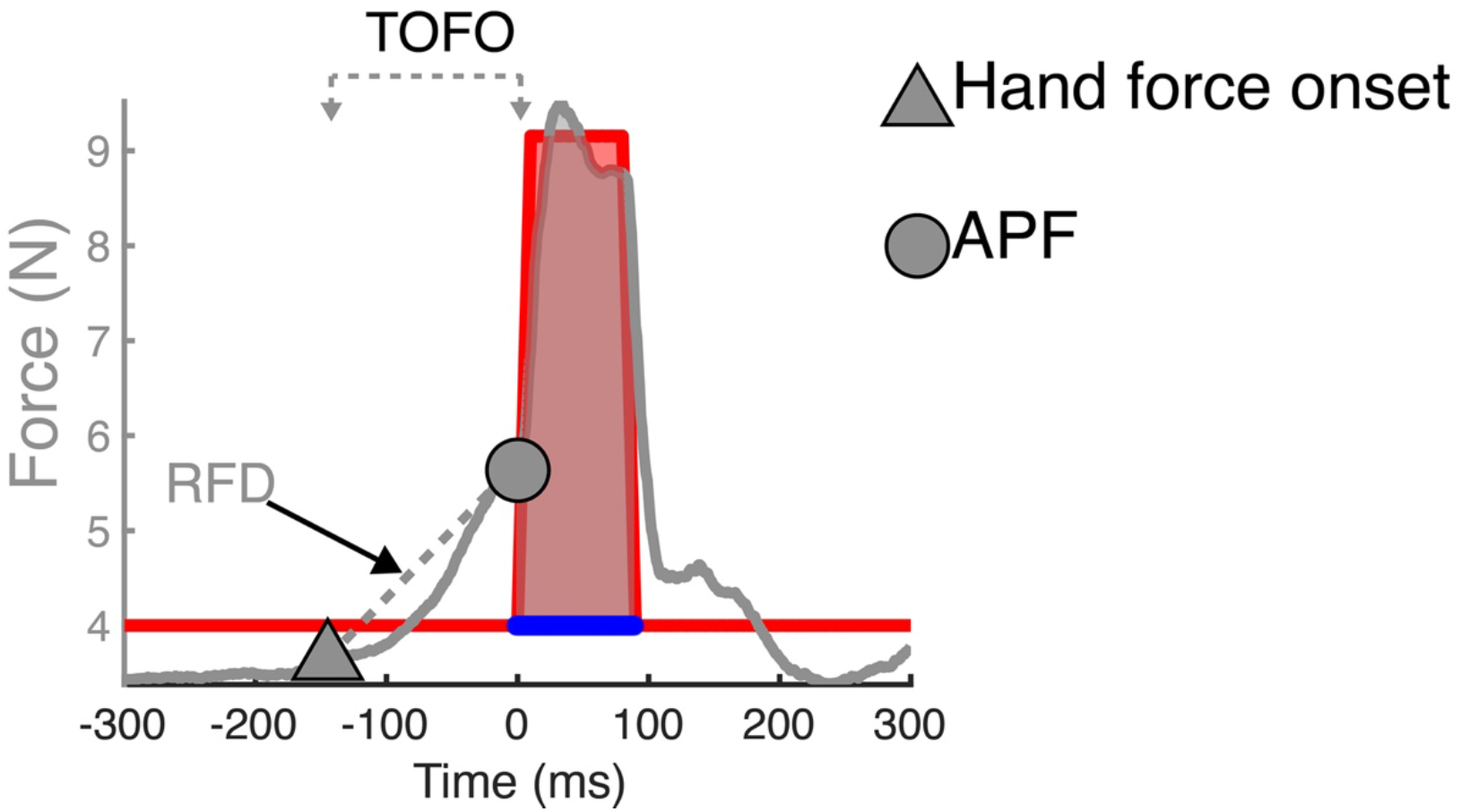
Performance variables for subsequent analysis. The y-axis shows the force applied by the participant in the Y direction (grey curve) to counter the impulsive force applied by the robot (red curve) during the contact with the virtual object. The shaded area indicates the impulse applied by the robot (red) and participant (grey). The triangle indicates the hand force onset. The circle indicates the anticipatory peak force (APF). Rate of force development (RFD) is the fitted slope between hand force onset and APF. Time of force onset (TOFO) is defined as the distance between the hand and object at hand force onset, divided by object speed.

We then calculated two parameters for the feedforward control of hand force. First, we calculated the peak hand force at contact (t = 0) and called it the *anticipatory peak force* (APF). We then fit a first order polynomial to the force data between the hand force at the time of hand force onset and APF and computed the slope of the fitted line. This slope provided a measure of rate of force development (RFD) between hand force onset and APF.

Finally, we calculated the distance along the Y-axis between the hand and the object at hand force onset. We then computed the *time of hand force onset*, TOFO, by dividing this variable by object speed (see Equation 3) (Lee 1998; Tresilian 1999).

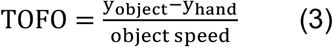

### Statistics

We conducted two-way repeated measures ANOVAs using object momentum (*Low* or *High*) and subcondition (*Mass* or *Speed*) as within-subject factors to compare collision performance, hand impulse, ΔImpulse, RFD, APF, and TOFO across conditions. The level of significance was set at a = 0.05 and effect sizes are reported using generalized η^2^. Post-hoc comparisons were performed using paired *t*-tests, with adjusted *p* values using the Bonferroni method. All values are reported as mean ± SE. All statistical analyses were performed in R (version 4.0.4).

## Results

### Limb force during collision depends on object momentum

Participants applied impulses within the acceptable range of 95-105% of the impulse applied by the robot in 48.07 ± 1.40%across all conditions. There was a small main effect of object momentum on the percentage of correct trials (*F*(1,19) = 5.35, *p* = 0.03, η^2^ = 0.01). Post-hoc comparisons showed participants performed better in the *Mass High* (51.3 ± 2.6%) than in the *Mass Low* (46.4 ± 2.96%) subcondition (*p* = 0.01), and no difference between the *Speed High* (47.7 ± 2.88%) and *Speed Low* (46.9 ± 2.83%) sub-conditions (*p* = 1.00). On trials that participants did not match the robot impulse correctly, they were more likely to apply too much force (overshoot) at the *Low* momentum conditions (main effect of momentum: *F*(1,19) = 48.91, *p* < 0.001, η^2^ = 0.16) and to not apply enough force (undershoot) in the *High* momentum conditions (main effect of momentum: *F*(1,19) = 24.03, *p* < 0.001, η^2^ = 0.08). Post-hoc comparisons showed that this pattern of overshooting and undershooting the ideal impulse was consistent for both *Mass* and *Speed* sub-conditions (all *p’s* < 0.05) (Fig. 4A).

**Figure 4.**
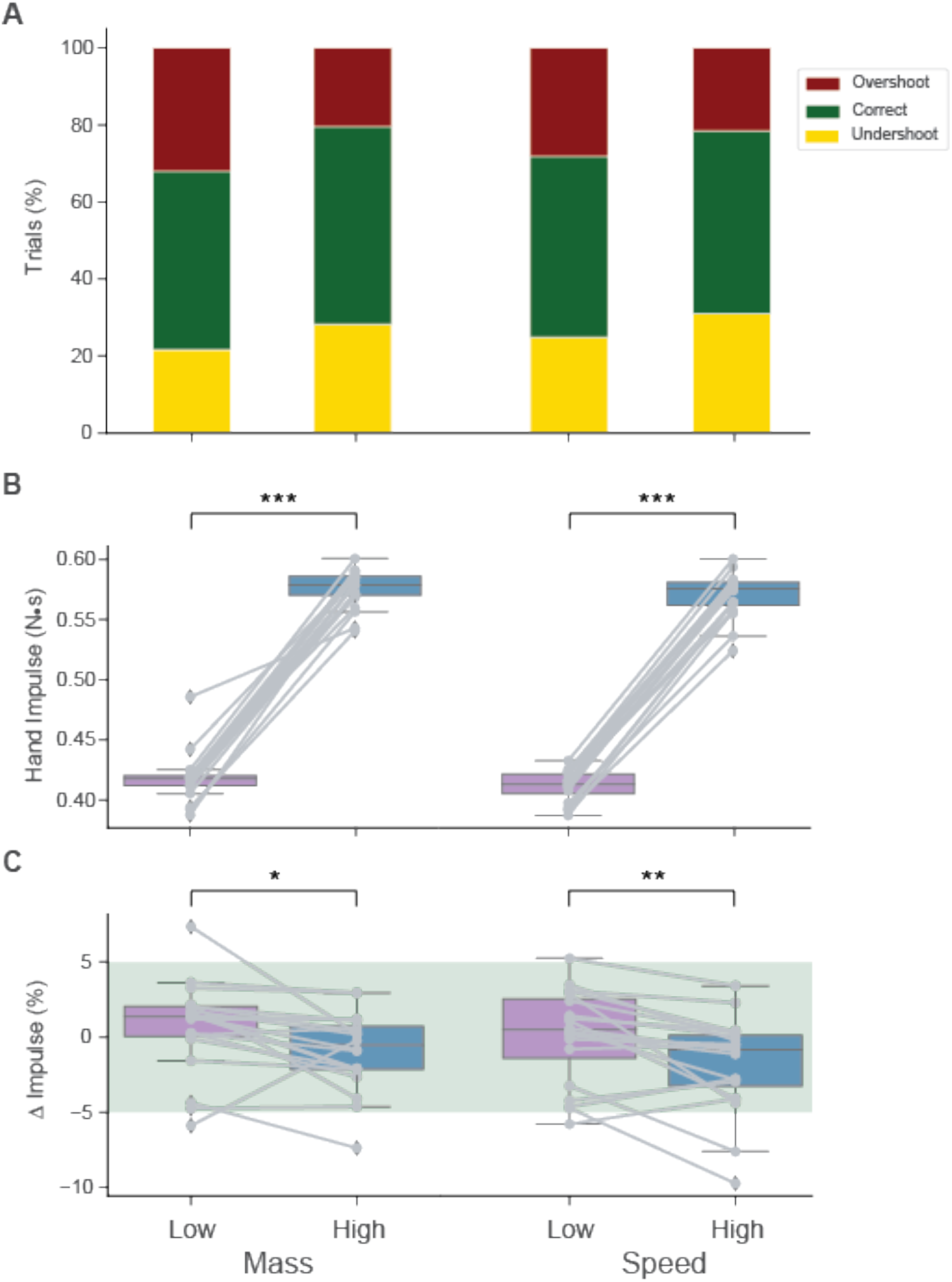
Limb force during collision. A) In unsuccessful trials participants tended to overshoot the hand impulse for Low momentum conditions, and to undershoot High momentum conditions. B) On average, participants’ hand impulse matched the robot’s impulse. C) Percentage difference between the impulse applied by the participant and the robot. The shaded area indicates the error margin allowable for a trial to be categorized as correct. Post-hoc differences (Bonferroni-adjusted): * indicates p < 0.05, ** indicates p < 0.01, *** indicates p < 0.001.

On average, participants were able to match the applied robot impulses of 0.41 N.s (*Low*) and 0.58 N.s (*High*) almost exactly: there was a large main effect of object momentum on the limb impulse (*F*(1,19) = 2980.05, *p* < 0.001, η^2^ = 0.96), with larger hand impulses for *Mass High* (0.58 ± 0.004 N.s) and *Speed High* (0.57 ± 0.004 N.s) than *Mass Low* (0.42 ± 0.005 N.s) and *Speed Low* (0.41 ± 0.003 N.s) *p’s* < 0.001). This confirms that the task was doable in that participants proportionately increased their hand impulse with an increase in object momentum due to increases in either speed or mass (Fig. 4B).

There was a significant main effect of object momentum on ΔImpulse (*F(1*,19) = 23.65, *p* < 0.001, η^2^ = 0.08). The average ΔImpulse was slightly positive in the *Mass Low* sub-condition (0.66 ± 0.68%), whereas it was significantly smaller and negative in the *Mass High* sub-condition (−0.92 ± 0.57%) (*p* = 0.03). Similarly, there was a significant difference in the average Δ Impulse between the *Speed Low* (0.09 ± 0.70%) and *Speed High* (−1.78 ± 0.70%) sub-conditions (*p* = 0.004), reflecting the tendency to overshoot the ideal force at lower momentum and undershoot the ideal force at higher momentum (Fig. 4C).

### Motor response amplitude and timing scales with object momentum

The rate of force increase between hand force onset and anticipatory peak force (APF) at collision, RFD, was higher in the *High* momentum conditions (main effect of momentum: *F(1*,19) = 82.58, *p* < 0.001, η^2^ = 0.12) for both mass (*Mass Low*: 27.4 ± 3.02 N/s; *Mass High:* 36.2 ± 3.54 N/s, *p* < 0.001) and speed (*Speed Low:* 26.3 ± 2.87 N/s; *Speed High:* 39.1 ± 3.87 N/s, *p* < 0.001) (Fig. 5A). There was a small but significant interaction such that the increase in RFD was larger when the increase in momentum was due to changes in speed than due to changes in mass (interaction: *F*(1,19) = 10.50, *p* = 0.004, η^2^ = 0.005). APF was also significantly higher in the *High* momentum conditions (main effect of momentum: *F*(1,19) = 247.45, *p* < 0.001, η^2^ = 0.15), for both changes due to mass (*Mass Low:* 6.15 ± 0.16 N; *Mass High:* 6.69 ± 0.16 N, *p* < 0.001) and speed (*Speed Low:* 6.16 ± 0.17 N; *Speed High:* 6.84 ± 0.19 N, *p* < 0.001) (Fig. 5B). However, in terms of absolute magnitude, this difference was rather small (9-11%) compared to the 41% increase in object momentum from the *Low* to the *High* momentum conditions (see Table 1).

**Figure 5.**
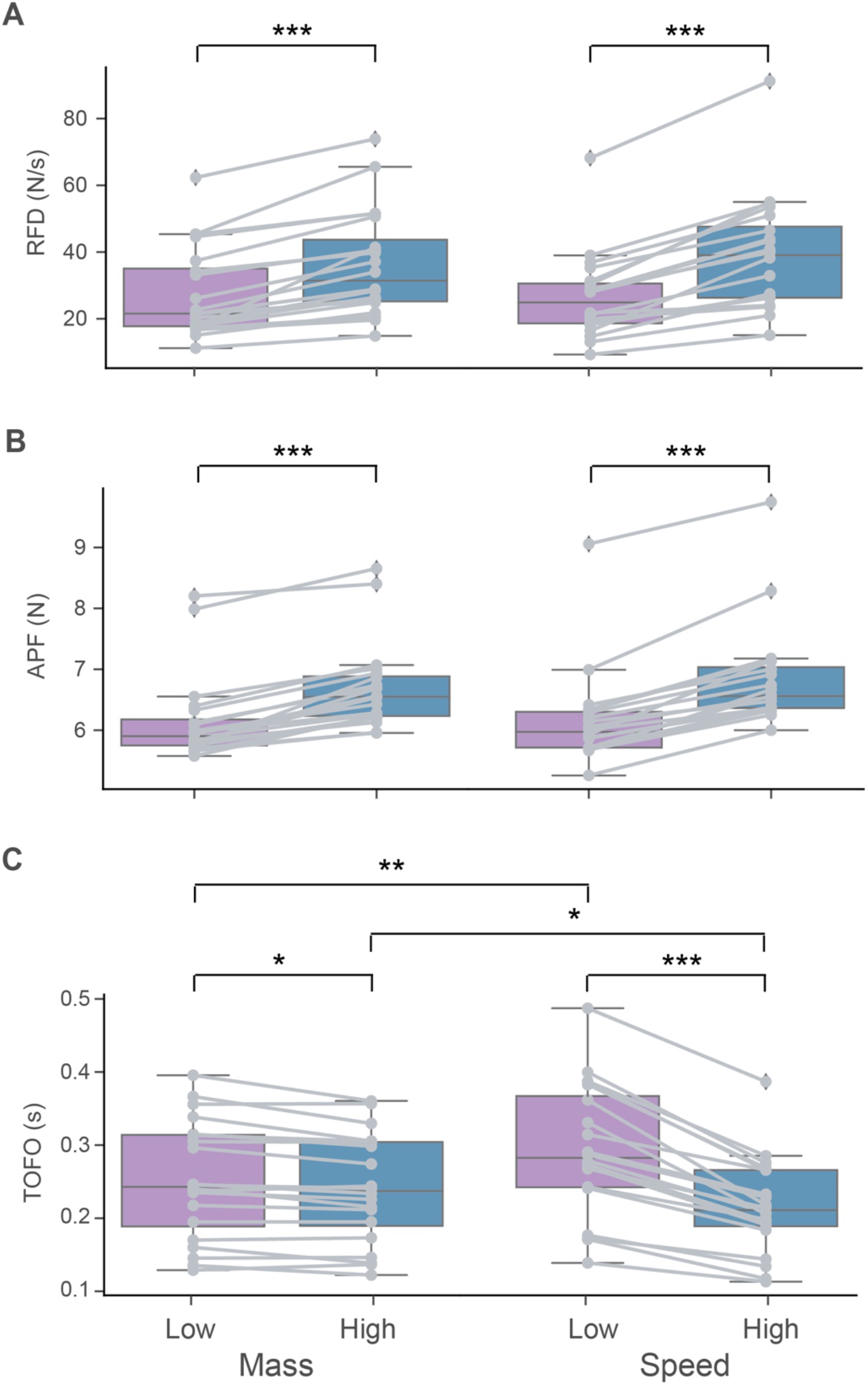
Motor response amplitude and timing. A) The rate of force development (RFD) scaled with object momentum. B) APF also scaled with momentum independently on how the momentum was manipulated. C) Hand force was increased above baseline levels closer to the time of contact in anticipation of higher momentum collisions. Post-hoc differences (Bonferroni-adjusted): * indicates p < 0.05, ** indicates p < 0.01, *** indicates p < 0.001.

There was a significant interaction between momentum and mass/speed sub-condition on the time of limb force onset, TOFO, (*F(1*,19) = 71.69, *p* < 0.001, η^2^ = 0.04). While TOFO decreased from *Low* to *High* momentum for both mass (*Mass Low*: 0.25 ± 0.02 s; *Mass High:* 0.24 ± 0.02 s, *p = 0.004*) and speed (*Speed Low:* 0.29 ± 0.02 s; *Speed High* 0.22 ± 0.02 s, *p* < 0.001), this decrease was much larger for the speed sub-conditions (Fig. 5C). Note that the average TOFO for both of the Mass sub-conditions was between the *Speed Low* and *Speed High* conditions, reflecting that the selected object speed for the Mass sub-conditions was an intermediate value (0.22 m/s) between Speed Low (0.18 m/s) and Speed (0.25 m/s) (see Table 1). Together, this suggests that in preparation for contact between the object and the hand, participants increased hand force above baseline levels much closer to the time of contact when anticipating a higher momentum collision, especially when the higher momentum was due to the object traveling at faster speeds. The distance of the target from the hand at which the hand force increased above baseline levels was similar across the different sub-conditions (*Mass Low:* 55 ± 4 mm; *Mass High:* 53 ± 4 mm; *Speed Low:* 53 ± 4 mm; *Speed High:* 53 ± 4 mm), suggesting that participants may have increased hand force when the target was at a certain distance from the hand. There was no main effect of momentum or any effect of the sub-condition.

## Discussion

In the current study, we introduced the *Mechanical STopping of Projectiles* (MSTOP) paradigm that simulated the mechanics of the interaction between the object and the hand based on Newton’s Second Law. This paradigm replicates the physics of the mechanical interaction between a projectile and the hand. We simulated the interaction such that the “momentum” (mass x velocity) of the object could be stopped by applying an equivalent mechanical impulse (integral area of force-time curve). If the applied impulse exceeded the object momentum, the object bounced back. If it was less than the object momentum, the object continued on its original trajectory. Note that the range of required performance was very strict, ±5% of the target, which is well below the standard Weber fractions for static force, ±10%, and led to many error trials, ~50%. This demanding paradigm was useful since it allows us to precisely control object dynamics. Unlike the real-world, where laws of motion exclusively dictate the mechanical interaction between projectiles and the arm, in augmented-reality we can decouple the physics of that interaction to systematically probe the relationship between object dynamics and limb motor control.

In different conditions, we either varied the speed or the mass of the object to alter its momentum. We hypothesized that visual system processes object momentum (mass x velocity) and feeds that information to the limb motor system to prepare postural responses. The prediction from this hypothesis was that the amplitude of the feedforward motor response would scale with the momentum of the object, regardless of whether the momentum increased due to speed or mass. Our results supported this prediction.

APF and RFD scaled with momentum in both the *Speed* and *Mass* sub-conditions - participants increased the hand force right before contact for objects with higher momentum, regardless of whether the momentum increased due to object speed or mass. This is consistent with the results of a previous study (Lacquaniti and Maioli 1989b).

We also predicted that the timing of the motor response initiation would be invariant to object motion and momentum. Our prediction was incorrect. We observed that when the object traveled at faster speeds, participants increased hand force closer to the time of contact. The distance of the object from the hand at the time of force increase was similar across all conditions, suggesting that participants may have initiated the force response when the objects were at a fixed distance. Previously, Port and colleagues showed that for objects moving at slow speeds, humans use a time-to-contact threshold to initiate hand movements, whereas for comparatively faster moving objects, they use a distance-to-contact threshold (Port et al. 1997). Our preliminary results support the distance-to-contact threshold in our augmented-reality paradigm. In the four experimental conditions, the mean values of the distance of the target at the point of hand force increase were within 2-3 mm of each other. The human eye can discern spatial distances separated by about 0.016° (1 arc minute). That would correspond to approximately 1.1 mm at a distance of 15 cm from the eye in the KINARM environment (30 cm depth). However, the magnitude of reaching errors made under memory-guided and open-loop (without visual feedback) conditions at that same distance tends to be ~4-10 mm (Heath et al. 2004). This suggests that the proprioceptive system could not have reliably differentiated the 2-3 mm difference that we saw across conditions. Thus, it seems very likely that participants initiated a force response when the object was at a fixed distance from the hand.

During many activities of daily living, humans interact with objects that are in relative motion with the body. These transient interactions require the limb to produce precise forces to stop the motion between the object and the body. Previously, researchers have used paradigms involving mechanical collisions between the hand and virtual objects (Bowman et al. 2009; White et al. 2011; Kuling et al. 2019) and hand and real projectiles (Lacquaniti and Maioli 1989b) to probe how the motor system stabilizes posture before and during collisions. In some of these paradigms, collisions have been modeled using Hooke’s Law assuming both the hand and the object are non-rigid. This is accomplished by programming a robot to apply position- (stiffness) and velocity-dependent (viscosity) forces on the limb during the collision.

A key feature of these existing paradigms is that collision between the hand and the object is primarily treated as a perturbation to the motor system to quantify pre-collision anticipatory and compensatory stretch-reflex like feedback responses during the collision. However, the mechanical interaction during the collision itself that dictates how force is transferred to projectiles from the hand has not to date been a focus of investigation. During this interaction, the limb produces a reactive force in response to the force generated by the object over the very short collision duration. Quantifying this interaction is important to better understand how humans acquire complex motor skills that require simultaneous force control and posture stabilization.

The present MSTOP paradigm has been designed to probe how humans control this force. We integrate the force applied over a short but modifiable time duration (fixed at 90 ms in this study) in realtime to calculate the net force impulse that is then transferred to the virtual object to change its momentum (Newton’s Second Law). More importantly, this paradigm allows us to manipulate object dynamics to systematically probe how humans learn and form novel sensorimotor maps to interact with moving objects. Future studies with this paradigm are well-positioned to address novel questions about control by manipulating object dynamics across trials and blocks to define how the sensorimotor maps between motion and force control are formed.

## Conclusion

In summary, we developed a new augmented-reality based mechanical stopping of projectiles (MSTOP) task which replicates the physics of a stopping task, but not the movement of the hands that are required to stop projectiles in the real-world. Using this paradigm, our study shows that object speed influences both the timing and amplitude of motor responses. We found that faster moving objects delayed the onset of the hand force response closer to the time of contact between the object and the hand. We also found that object momentum scaled the amplitude of the feedforward force response regardless of whether the momentum was increased due to speed or mass. Our results suggest that the timing and amplitude of hand motor responses in this virtual catching task are affected by different features of object motion. Future experiments using this paradigm will help us understand the mechanisms underlying these sensorimotor processes.

## Author contributions

AGG and TS conceived and designed research; AGG and SG performed experiments; AGG, DAB, and TS analyzed data; AGG, DAB, IK and TS interpreted results of experiments; AGG and TS prepared figures; TS, AG, IK, DAB drafted manuscript; AGG, IK, DAB, and TS edited and revised manuscript; AGG, IK, SG, DAB, and TS approved the final version of manuscript.

## Data Availability

The datasets generated during and/or analyzed during the current study are available from the corresponding author on reasonable request.

## References

Barany DA, Gómez-Granados A, Schrayer M, Cutts SA, Singh T (2020) Perceptual decisions about object shape bias visuomotor coordination during rapid interception movements. Journal of Neurophysiology 123:2235–2248 doi: 10.1152/jn.00098.2020

Bowman MC, Johannson RS, Flanagan JR (2009) Eye–hand coordination in a sequential target contact task. Experimental Brain Research 195:273–283 doi: 10.1007/s00221-009-1781-x

Brenner E, Smeets JB (2011) Continuous visual control of interception. Human Movement Science 30:475–494

Burdet E, Franklin DW, Milner TE (2013) Human Robotics: Neuromechanics and Motor Control. The MIT Press, Cambridge, MA

Burkholder TJ (2016) Model-based approaches to understanding musculoskeletal filtering of neural signals. In: Prilutsky BI, Edwards DH (eds) Neuromechanical Modeling of Posture and Locomotion. Springer New York, New York, NY, pp 103–120

Debats NB, Kingma I, Beek PJ, Smeets JB (2012) Moving the Weber fraction: the perceptual precision for moment of inertia increases with exploration force. PLoS One 7:e42941

Heath M, Westwood DA, Binsted G (2004) The control of memory-guided reaching movements in peripersonal space. Motor Control 8:76–106

Jones LA (1986) Perception of force and weight: theory and research. Psychological Bulletin 100:29–42

Kuling IA, Salmen F, Lefèvre P (2019) Grip force preparation for collisions. Experimental Brain Research 237:2585–2594 doi: 10.1007/s00221-019-05606-y

Kurtzer IL (2015) Long-latency reflexes account for limb biomechanics through several supraspinal pathways. Frontiers in Integrative Neuroscience 8

Lacquaniti F, Borghese N, Carrozzo M (1991) Transient reversal of the stretch reflex in human arm muscles. Journal of Neurophysiology 66:939–954

Lacquaniti F, Borghese N, Carrozzo M (1992) Internal models of limb geometry in the control of hand compliance. Journal of Neuroscience 12:1750–1762

Lacquaniti F, Carrozzo M, Borghese NA (1993) Time-varying mechanical behavior of multijointed arm in man. Journal of Neurophysiology 69:1443–1464

Lacquaniti F, Maioli C (1989a) Adaptation to suppression of visual information during catching. Journal of Neuroscience 9:149–159 doi: 10.1523/jneurosci.09-01-00149.1989

Lacquaniti F, Maioli C (1989b) The role of preparation in tuning anticipatory and reflex responses during catching. Journal of Neuroscience 9:134–148 doi: 10.1523/jneurosci.09-01-00134.1989

Lee DN (1976) A theory of visual control of braking based on information about time-to-collision. Perception 5:437–459

Lee DN (1998) Guiding movement by coupling taus. Ecological Psychology 10:221–250

Mrotek LA, Soechting JF (2007) Target interception: hand–eye coordination and strategies. Journal of Neuroscience 27:7297–7309

Port NL, Lee D, Dassonville P, Georgopoulos AP (1997) Manual interception of moving targets I. Performance and movement initiation. Experimental Brain Research 116:406–420 doi: 10.1007/pl00005769

Scott SH (2012) The computational and neural basis of voluntary motor control and planning. Trends in Cognitive Sciences 16:541–549

Singh T, Fridriksson J, Perry CM, Tryon SC, Ross A, Fritz SL, Herter TM (2017) A novel computational model to probe visual search deficits during motor performance. Journal of Neurophysiology 117:79–92 doi: https://doi.org/10.1152/jn.00561.2016

Tresilian JR (1991) Empirical and theoretical issues in the perception of time to contact. Journal of Experimental Psychology: Human Perception and Performance 17:865

Tresilian JR (1999) Visually timed action: time-out for ‘tau’? Trends in Cognitive Sciences 3:301–310

Tresilian JR (2005) Hitting a moving target: perception and action in the timing of rapid interceptions. Perception & Psychophysics 67:129–149

White O, Thonnard J-L, Wing A, Bracewell R, Diedrichsen J, Lefevre P (2011) Grip force regulates hand impedance to optimize object stability in high impact loads. Neuroscience 189:269–276

Zago M, Lacquaniti F (2005) Cognitive, perceptual and action-oriented representations of falling objects. Neuropsychologia 43:178–188

